# Lifespan analysis of dystrophic *mdx* fast-twitch muscle morphology and its impact on contractile function

**DOI:** 10.1101/2021.09.07.459226

**Authors:** Leonit Kiriaev, Sindy Kueh, John W. Morley, Kathryn N. North, Peter J. Houweling, Stewart I. Head

## Abstract

Duchenne muscular dystrophy is caused by the absence of the protein dystrophin from skeletal muscle and is characterized by progressive cycles of necrosis/regeneration. Using the dystrophin deficient *mdx* mouse model we studied the morphological and contractile chronology of dystrophic skeletal muscle pathology in fast twitch EDL muscles from animals 4-22 months of age containing 100% regenerated muscle fibers. Catastrophically, the older age groups lost ∼80% of their maximum force after one eccentric contraction of 20% strain, with the greatest loss ∼93% recorded in senescent 22 month old *mdx* mice. In old age groups there was minimal force recovery ∼24% after 120 minutes, correlated with a dramatic increase in the number and complexity of branched fibers. This data supports our two-stage model where a “tipping point” is reached when branched fibers rupture irrevocably on eccentric contraction. These findings have important implications for pre-clinical drug studies and genetic rescue strategies.

## INTRODUCTION

In Duchenne Muscular dystrophy (DMD), the absence of dystrophin from skeletal muscle triggers necrosis. The initial wave of necrosis is followed by regeneration of the skeletal muscle tissue and subsequent cycles of necrosis and regeneration ^1, 2^. The same pathology occurs in the most commonly used animal model of DMD, the dystrophin-deficient *mdx* mouse, where the long form of dystrophin is absent from the inner surface of the skeletal muscle sarcolemma ^3–7^. In the *mdx* mouse many studies have shown that the predominantly fast-twitch Extensor Digitorum Longus (EDL) and Tibialis Anterior (TA) muscles are more susceptible to eccentric contraction (EC) induced force deficit compared with dystrophin positive controls ^8–12^. It is interesting to note that in the dystrophin deficient *mdx* mouse muscles which lack type 2B (fast-twitch glycolytic) fibers (such as the soleus) are not susceptible to EC force deficits, even though the absence of dystrophin in these muscles triggers waves of necrosis and regeneration such that the muscle fibers are completely replaced by 8 weeks of age ^13^. The EC induced drop in absolute force in fast-twitch muscles has been widely attributed to being the result of sarcolemmal damage and is commonly used as a model of membrane damage in the dystrophinopathies ^8, 9, 12, 14–26^. Studies designed to investigate possible treatments or cures for DMD commonly utilize the *mdx* mouse as a preclinical model to examine if the pharmaceutical/genetic intervention strategies prevents or reduces this EC force loss in dystrophin deficient fast-twitch skeletal muscle ^27–35^. We believe these studies are flawed in cases where they assume the EC force deficit is due to sarcolemmal rupture or tearing because there is a compelling body of evidence ^36–42^ demonstrating that in younger *mdx* mice, this EC deficit in the fast-twitch muscles is prevented by molecules that block stretch sensitive ion channels and also by exposing the dystrophic muscle to antioxidants before the EC ^43^. Recent support of this non-sarcolemmal damage pathway came from the work of Olthoff, et al. ^44^ where they demonstrated that EC force deficit could largely be reversed if the dystrophin deficient fast-twitch muscle was allowed to recover for a period of up to 120 minutes or if the period between repeated ECs was lengthened from the commonly used 3-5 minutes to 30 minutes. Additionally, Olthoff, et al. ^44^ showed that EC force deficit in younger 3 month old *mdx* fast-twitch muscles was reduced by bath application of the antioxidant N-Acetyl Cysteine (NAC) or by genetically upregulating an endogenous antioxidant. This adds further weight to Allen’s proposal (see Allen, et al. ^36^ for a review) that in dystrophinopathies, muscles have a greater susceptibility to ROS-induced Ca^2+^ influx via abnormally functioning ion channels ^40, 41^ and it is the elevated [Ca^2+^]in that triggers the cycles of necrosis/regeneration in dystrophinopathies. Fast-twitch muscles from *mdx* mice lose up to 90% of their initial force generating capacity depending upon the EC protocol used (normally a series of between 3-120 contractions of varying length and velocity)^45^. During the course of a series of ECs the force deficit occurs incrementally with each EC, rather than as an abrupt drop in force on the first EC which would be predicted if the sarcolemma was tearing or ripping. The exception to this is when you look at fast-twitch muscles from older (58-112 weeks) *mdx* mice where most of the force loss occurs abruptly during the first EC ^12^. It is important to note that this is not an ageing effect, as a similar phenomenon is not seen in the age matched littermate controls. We and others have shown that throughout the 30% shorter ^46, 47^ lifespan of the *mdx* mouse, skeletal muscle undergoes continuous cycles of degeneration and regeneration ^6, 19–26, 46, 48, 49^.

Regenerated fibers are characterized by having central nuclei and as the number of cycles increases so do the incidences of abnormal branched morphology ^8, 12, 25, 50–55^. As the *mdx* mouse ages the branched fibers become more chaotic and bizarre in appearance, with a single continuous cytoplasmic syncytium being capable of supporting ten or more major branches ^8, 12, 25^. Once the number of branched fibers and their complexity of branching exceeds a threshold we refer to as “tipping point”, we and others have hypothesized that it is the branching, in and of itself, that weakens the muscle ^12, 15, 16, 49, 52, 56–59^. Previously our laboratory has shown this by looking at EDL muscles from the *mdx* dystrophic mouse from 2 weeks up to 112 weeks of age and it’s only in the muscles greater than 6 months of age that there is a sudden loss of force with the first EC. Subsequent imaging of the muscle fibers *in–situ* and also examination of enzymatically liberated single fibers show multiple examples of breaking and rupturing at branch points. In the present study we use a strong EC protocol which produces force deficits in littermate control and *mdx* mice in adolescent to senescent aged groups and allow the muscles to recover for 120 minutes post ECs to see if there is a non-recoverable force deficit which is correlated with fiber branching in dystrophic muscles.

## METHODS

### Ethics approval

Animal use was approved by the Western Sydney University Animal Care and Ethics Committee. A12907. These experiments were conducted in compliance with the animal ethics checklist and ethical principles under which the journal operates.

### Animals

The majority of previous dystrophic muscle function studies used separate colonies of wild-type control and dystrophic mice which have been inbred for over 25 years introducing the possibility of new mutations to the groups.

In this study, littermates are bred to act as control animals for dystrophic mice. These are genetically more appropriate controls for dystrophin studies as both dystrophin negative and positive animals are from identical genetic backgrounds ^60^.

Male dystrophic mice with littermate controls were obtained from the Western Sydney University animal facility. The colony of dystrophic mice & littermate controls used in this study were second generation offspring crossed between female C57BL/10ScSn-Dmd^m^*^dx^* and male C57BL/10ScSn mice. Littermate controls were distinguished from dystrophic mice by genotyping.

Mice from five age groups were used in this study: 4, 9, 15, 18 and 22 months old where dystrophic muscles have undergone at least one round of necrosis/regeneration ^13^. These age groups were selected to breach the gap in dystrophic literature investigating muscle performance and recovery from contraction induced damage in adolescent to senescent mice. They were housed at a maximum of four to a cage in an environment controlled room with a 12h-12h light-dark cycle. Standard rodent pellet chow (Gordon’s Specialty Stockfeeds, Yanderra, NSW, Australia) and water were available ad libitum. Basic enrichments such as nesting crinkle material and polyvinyl chloride pipe tube were provided.

A total of 71 male mice were used in this study (39 *mdx* mice and 32 littermate control mice), 135 Extensor digitorum longus (EDL) and 117 Tibialis anterior (TA) muscles collected for this study, not all of which were used in every procedure.

### Muscle preparation

Mice were placed in an induction chamber and overdosed with isoflurane delivered at 4% in oxygen from a precision vaporizer. Animals were removed when they were not breathing and a cervical dislocation was immediately carried out. Both the fast-twitch EDL and TA muscles were dissected from the hind limb which was submerged in oxygenated Krebs solution at all times during the dissection. TA muscles were dissected tendon to tendon, trimmed (excess tendons) and weighed. The dissected EDL muscle was tied by its tendons from one end to a dual force transducer/linear tissue puller (300 Muscle Lever; Aurora Scientific Instruments, Canada) and secured to a base at the other end using 6-0 silk sutures (Pearsalls Ltd, UK).

Each muscle was then placed in a bath containing Krebs solution (also used as dissection solution) with composition (in mM): 4.75 KCl, 118 NaCl, 1.18 KH_2_PO_4_, 1.18 MgSO_4_, 24.8 NaHCO_3_, 2.5 CaCl_2_, 10 glucose, 0.1% fetal calf serum and bubbled continuously with carbogen (95% O_2_, 5% CO_2_) to maintain pH at 7.4. The muscle was stimulated by delivering a current between two parallel platinum electrodes, using an electrical stimulator (701C stimulator; Aurora Scientific Instruments). All contractile procedures were designed, measured and analyzed using the 615A Dynamic Muscle Control and Analysis software (Aurora Scientific Instruments). At the start of each experiment, the muscle was set to optimal length (Lo) which produces maximal twitch force. Muscle dissection and experiments were conducted at room temperature (∼20-22°C).

### Initial maximum force and contractile protocols

An initial supramaximal stimulus was given at 125Hz (1ms pulses) for 1s and force produced recorded as Po, the maximum force output of the muscle at L_o_. Pairs of EDL muscles from each animal were divided into two test groups that undergo either force frequency or isometric contraction protocols. Immediately after, all muscles were left to rest for 5 minutes during which muscle length was reset to L_o_, followed by the EC and recovery procedures.

### Force frequency curve

Force-frequency curves were generated for one set of muscles to measure contractile function. Trains of 1ms pulse stimuli performed at different frequencies (2, 15, 25, 37.5, 50, 75, 100, 125, 150Hz) were given for 1s, the force produced measured and a 30s rest was allowed between each frequency. A sigmoid curve relating the muscle force (P) to the stimulation frequency (f) was fitted by linear regression to this data.

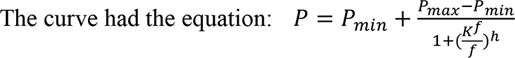

From the fitted parameters of the curve, the following contractile properties were obtained: force developed at minimal (P_min_) and maximal (P_max_) stimulation at the conclusion of the force-frequency curve. Half-frequency (K_f_) is the frequency at which the force developed is halfway between (P_min_) and (P_max_), and Hill coefficient (h) which quantifies the slope of the muscle force frequency sigmoidal curve. These were used for population statistics.

### Isometric contractions

The contralateral EDL muscle was subjected to 10 consecutive isometric (fixed-length) supramaximal tetanic contractions, each lasting 1s separated by a 60s rest, then given a recovery contraction 5 minutes after the protocol (total of 11 contraction). Stimulus pulses within the 1s tetanus were 0.5ms in duration (width) and were delivered at a frequency of 125Hz. This protocol is a modified version of the sequence used by Claflin and Brooks ^61^. The force measured at each isometric contraction was expressed as a percentage of the force produced during the first (Initial) contraction.

### Eccentric contractions and recovery

A series of eccentric (lengthening) contractions were then performed on each EDL where the contracted muscle was stretched 20% from L_o_. At t=0s, the muscle was stimulated via supramaximal pulses of 1ms duration and 125Hz frequency. At t=0.9s, after maximal isometric force was attained, each muscle was stretched 20% longer than their optimal length and held at this length for 2s before returning to L_o_. Electrical stimulus was stopped at t=5s. The EC procedure was repeated 6 times with 3 minute rest intervals. This is followed immediately by a recovery protocol. The force measured at each EC was expressed as a percentage of the force produced during the first (Initial) contraction. From the first EC data trace, baseline force was taken before and after the first EC to quantify baseline changes as a result of the first EC.

Upon completion of the 6 EC contraction sequence, recovery force was measured immediately (Post 0’) through isometric contractions given at 125Hz (1ms pulses) for 1s as well as at the 20, 40, 60, 80, 100 and 120 minute time points (Post 20’, Post 40’, etc.). These force values were then expressed as a percentage of the Po measured before the 6 ECs. The recovery protocol is a modified version of the sequence used by Olthoff, et al. ^44^.

### Twitch kinetics

Twitch kinetics was measured at 3 time points throughout the contractile protocol to compared differences between (1) pre EC, (2) post EC and (3) post recovery kinetics. The twitches were performed at 2Hz, for 1ms (width) and 1s duration immediately after initial maximum force (Pre), after the EC protocol (Post) and recovery protocol (Recov). The following parameters were collected: twitch force, half relaxation time (HRT) and time to peak (TTP).

### Muscle Stiffness

Stiffness is the resistance of an elastic body to deflection or deformation by an applied force. As an indicator of active muscle stiffness (during contraction), the change in force as the muscle is lengthened was measured during the first EC. Force was expressed as a percentage of the isometric force before stretching, and length was expressed as a percentage of optimum length. To estimate muscle stiffness, the change in muscle force was divided by the change in muscle length (as a percentage of L_o_) during the first EC.

Stiffness is an inverse indication of muscle compliance; a stiffer muscle would exhibit a greater change in force for a given change in length. Earlier skinned fiber studies of *mdx* single muscle fibers where all extracellular components were removed reported no major differences in the function of contractile proteins ^25^. Thus investigating stiffness provides an indication of the effects of non-contractile elements on muscle function.

### Work done

Work is an energy quantity given by force times distance, this measurement was used to provide a quantitative estimate of the eccentric damage-inducing forces. Work done to stretch the muscle was calculated from the force tracings through multiplying the area underneath the lengthening phase of the force tracing with the velocity of lengthening.

Several studies have examined the effect of various parameters on the force deficit produced by ECs, and found that the work done to stretch the muscle during the lengthening phase of an EC was the best predictor of the magnitude of the force deficit ^45, 62–64^.

### Muscle mass and cross sectional area

After contractile procedures were completed, the EDL muscle was removed from the organ bath and tendons trimmed. Both the EDL and TA muscles were blotted lightly (Whatmans filter paper DE81 grade) and weighed using an analytical balance (GR Series analytical electronic balance).

Physiological cross sectional area (PCSA) was calculated through dividing the muscle mass by the product of its length and mammalian muscle density. Specific muscle force was obtained through dividing raw force values by cross sectional area. When normalizing force using a calculation of physiological cross sectional area in mouse EDL many studies use a correction factor to allow for fiber length (Lf). We chose not to in the present study because:

i. There is considerable variation in the literature as to the value of Lf/Lo in mouse EDL muscle with a ratios of 0.44, 0.68, 0.75 and 0.85 being used ^65–68^. Depending on which one is adopted, widely varying values for specific force will be obtained. In this study we were not primarily concerned with the actual values of the specific force; we were mainly interested in seeing whether it differed between the ages and genotype.
ii. As we show here branched fibers are found in the dystrophic muscles, and the measurement of fiber length becomes especially problematic in branched fibers. The application of a uniform Lf/Lo across the whole muscle might not be valid due to the different geometry of branched fibers and unbranched fibers. It is not clear how the length of the fiber would be defined. It is possible that some other method of estimating total fiber cross sectional area (CSA) is necessary in a muscle containing branched fibers.

However, we did normalize forces with respect to an estimate of physiological cross-sectional area, according to the equation CSA=MM/(L_o_*D), where MM is the muscle mass, L_o_ is the optimal length and D is the density of skeletal muscle (1.06 g/cm^3^), to enable us to compare muscle of differing sizes and weights ^69^. In healthy rodent hind limb muscle maximal tension was found to be directly proportional to calculated PCSA ^70^. However, this method still not ideal as we are assuming that the muscle density is unaltered by fat and connective tissue infiltration which is particularly problematic in the old dystrophic animals, shown to have extensive accumulation of connective tissue and some fat infiltration ^71^.

### Skeletal Muscle single fiber enzymatic isolation and morphology

Following contractile procedures and weighing, EDL muscles were digested in Krebs solution (without FCS) containing 3 mg/ml collagenase type IV A (Sigma Aldrich, USA), gently bubbled with carbogen (95% O_2_, 5% CO_2_) and maintained at 37°C. After 25 minutes the muscle was removed from solution, rinsed in Krebs solution containing 0.1% fetal calf serum and placed in a relaxing solution with the following composition (mM): 117 K^+^, 36 Na^+^, 1 Mg^2+^, 60 Hepes, 8 ATP, 50 EGTA (Note: internal solution due to chemically skinning by high EGTA concentration). Each muscle was then gently agitated using pipette suction, releasing individual fibers from the muscle mass. Using a pipette 0.5ml of solution was drawn and placed on a glass slide for examination and photographs of dissociated fibers taken. In instances where a long fiber covered several fields of the microscope view a series of overlapping photomicrographs were taken and these were then stitched together using the Coral draw graphic package.

A total of 11657 fibers from 65 EDL muscles were counted; 7119 fibers from 34 controls and 4538 from 31 dystrophic muscles. Only intact fibers with no evidence of digestion damage were selected for counting.

### Statistical analyses

Data was presented as means ± SD, Differences occurring between genotypes and age groups were assessed by two-way ANOVA, genotype being one fixed effect and age groups the other, their interactions included. *Post hoc* analysis was performed using Sidak’s multiple comparisons test. All tests were conducted at a significance level of 5%. All statistical tests and curve fitting were performed using a statistical software package Prism Version 7 (GraphPad, CA, USA).

## RESULTS

### Degree of fiber branching and complexity with age

The number of branched fibers and complexity of fiber branching within a single branched fiber syncytium found in *mdx* EDL muscles is shown in Figure 1 (note these counts represent all fibers counted and there are no error bars). The evidence ^8, 12, 13, 15, 25, 51–56, 72–74^ now overwhelming supports the findings shown in Figure 1, that as the *mdx* animals age the number and complexity of branched fibers increases dramatically. Single enzymatically isolated EDL fibers from littermate control animals showed between <1% to ∼3% branching in all age groups (consisted with reports from other studies)^8, 12, 16, 25, 51, 55, 73, 75, 76^. In the 4 month *mdx* 48% of all fibers counted are branched; the branching is relatively simple with 41% of branched fibers having one or two branches per fiber. By the time they reach 9 months of age 83% of fibers contain branches, of these 38% of fibers contain one or two branches and 45% fibers with 3 or more branches per fiber syncytium. At 15 and 18 months of age dystrophic EDL muscles have 96% branched fibers of which 64% have 3 or more branches per fiber syncytium and 32% having one or two branches. At 22 months of age the majority of *mdx* EDL fibers contained branches with 77% showing 3 or more branches per fiber.

**Figure 1:**
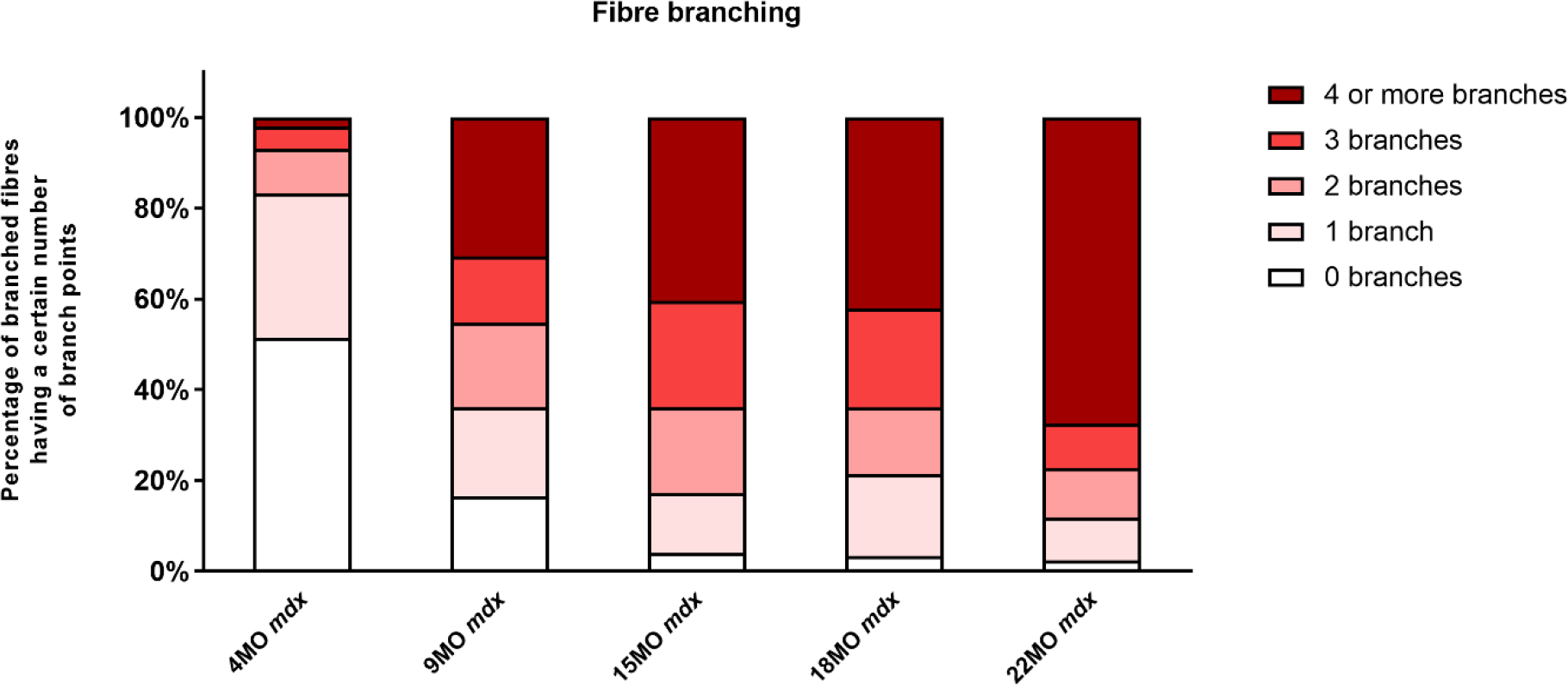
The degree of fiber branching in EDL muscles as a percentage of all fibers by categorizing them based on number of branch points. Fibers from littermate controls were omitted due to very low presence of fiber branching, less than 1%. Note this is a bar graph of all fibers counted and as such is an absolute measure, all photomicrographs were taken from these fiber counts. A total of 11657 fibers from 65 EDL muscles were counted; 7119 fibers from 34 controls and 4538 fibers from 31 *mdx* muscles. For 4 month group: n=1514 control n=1030 *mdx* fibers, 9 month group: n=1320 control n=1130 *mdx* fibers, 15 month group: n=1615 control n=1263 *mdx* fibers, 18 month group: n=1435 control n=605 *mdx* fibers and 22 month group: n=1235 control n=511 *mdx* fibers.

### Mass, length, and cross sectional area

TA muscles were heavier for *mdx* animals compared to littermate controls at all age groups (Figure 2a), peak differences in muscle mass occurring at 15 months (MD 37.3, 95% CI [26.39, 48.21], P<0.0001) compared to 4 months (MD 18.34, 95% CI [8.92, 27.76], P<0.0001) and 22 months (MD 19.12, 95% CI [5.65, 32.58], P=0.0016) age groups. These findings and a lack of age effects on muscle mass are consistent with dystrophic TA studies published previously ^10, 19, 71, 77, 78^. The same increase in dystrophic muscle mass can also be seen in the EDL (Figure 2b), peak differences likewise occurring at 15 months of age (MD 8.52, 95% CI [6.09, 10.96], P<0.0001) compared to 4 month (MD 3.18, 95% CI [0.74, 5.62], P=0.0045) and 22 month (MD 6.75, 95% CI [4.08, 9.43], P<0.0001) age groups. Muscle hypertrophy in *mdx* muscle is a recognized feature ^12, 19, 20, 22–24, 48, 69, 79, 80^ and can largely be attributed to the fiber branching which occurs the regenerated dystrophic fibers ^8, 12, 55, 74^. It should be noted that both TA & EDL control muscle mass has remained consistent with age and there was no evidence of sarcopenia up to 22 months of age. Other properties for the EDL such as length and physiological cross sectional area for all age groups are shown in Table 1. EDL muscle length remained the same regardless of genotype or age, hence when physiological cross sectional area was calculated for each muscle the differences can be attributed to those seen in EDL muscle mass.

**Figure 2:**
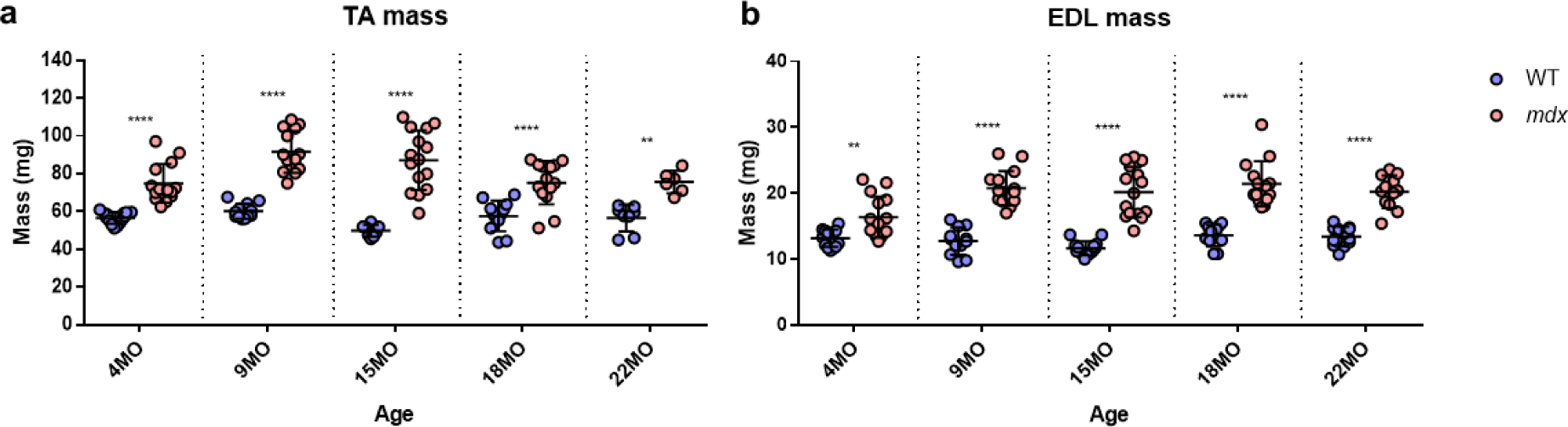
Muscle mass for all age groups. **a)** Scatterplot of TA muscle mass for 4 month (n=14 control; n=14 *mdx*), 9 month (n=12 control; n=14 *mdx*), 15 month (n=8 control; n=15 *mdx*), 18 month (n=12 control; n=14 *mdx*) and 22 month (n=8 control; n=6 *mdx*) age groups. **b)** Scatterplots of EDL muscle mass for 4 month (n=14 control; n=15 *mdx*), 9 month (n=12 control; n=15 *mdx*), 15 month (n=14 control; n=15 *mdx*), 18 month (n=12 control; n=14 *mdx*) and 22 month (n=12 control; n=12 *mdx*) age groups. In each group the horizontal line indicates the mean value ± SD. There were no statistical differences between age groups, statistical differences displayed within graphs are differences between genotypes assessed by two-way ANOVA, *post hoc* analysis using Sidak’s multiple comparisons test. ****P<0.0001 and **0.001<P<0.01.

**Table 1:**
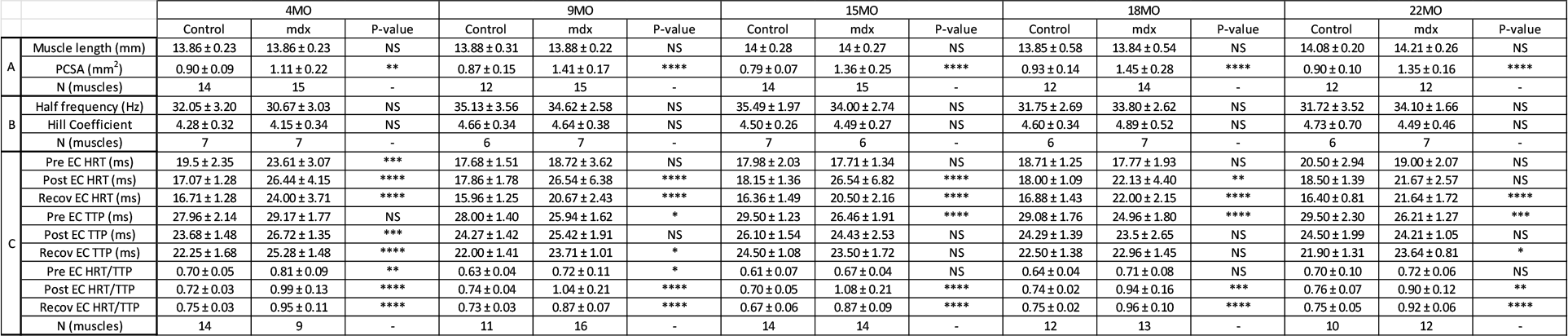
Statistical analyses and sample size of muscle properties, force frequency parameters and kinetics for EDL muscles across all age groups. . Muscle properties *(Row section 1*): Muscle length and calculated physiological cross sectional area. Force frequency parameters *(Row section 2*): Half frequency and Hill coefficient. Kinetics *(Row section 3*): Half relaxation time (HRT), Time to peak (TTP) and HRT/TTP ratio. Twitch kinetics were measured (1) pre-EC, (2) post-EC and (3) after recovery from the EC protocol. All data shown are mean ± SD. No statistical differences were found between age groups, P-values show genotype effects assessed by two-way ANOVA, *post hoc* analysis using Sidak’s multiple comparisons test. ****P<0.0001, ***0.0001<P<0.001, **0.001<P<0.01, *0.01<P<0.05 and NS labelled for no statistical significance.

### Maximal tetanic and twitch force

The EDL muscle isometric maximum force production (P0) for all age groups are presented in Figure 3a. Four month old mice showed no significant differences in force production between dystrophic and littermate control muscles, however, as the animals age 9-22 months, the maximum force generated by the dystrophic EDL is significantly less than age matched littermate controls. The low force output is most pronounced in older mice, where 18 month *mdx* muscles produced ∼25% less force than controls (MD -92.95, 95% CI [-146.2, -39.73], P<0.0001) and 22 month *mdx* muscles produced ∼57% less force than controls (MD -197.1, 95% CI [-256.2, -138], P<0.0001). Interestingly, the 22 month old dystrophic cohort produced significantly less force compared to all other age groups (F (4, 111)=9.26, P<0.0001). When each EDL muscle’s force was corrected for physiological cross sectional area (Figure 3b) the 22 month old dystrophic cohort still produced significantly less specific force compared to dystrophic muscles from all other age groups (F (4, 107)=7.85, P<0.0001). Most notably dystrophic EDL muscles produced significantly less specific force compared to littermate control animals at all age groups. At 4 months of age the *mdx* muscles produce ∼20% less force c.f. controls (MD -84.92, 95% CI [-156.6, -13.28], P=0.012), this deficit increases at 15 months with *mdx* muscles producing ∼54% less force c.f. controls (MD - 262.5, 95% CI [-331.9, -193], P<0.0001) and peaks at 22 months of age where dystrophic EDL muscles generating ∼73% less force than controls (MD -279.4, 95% CI [-352.7, -206.1], P<0.0001) which correlated with the increase in the number and complexity of branched fibers during this period Figure 1.

**Figure 3:**
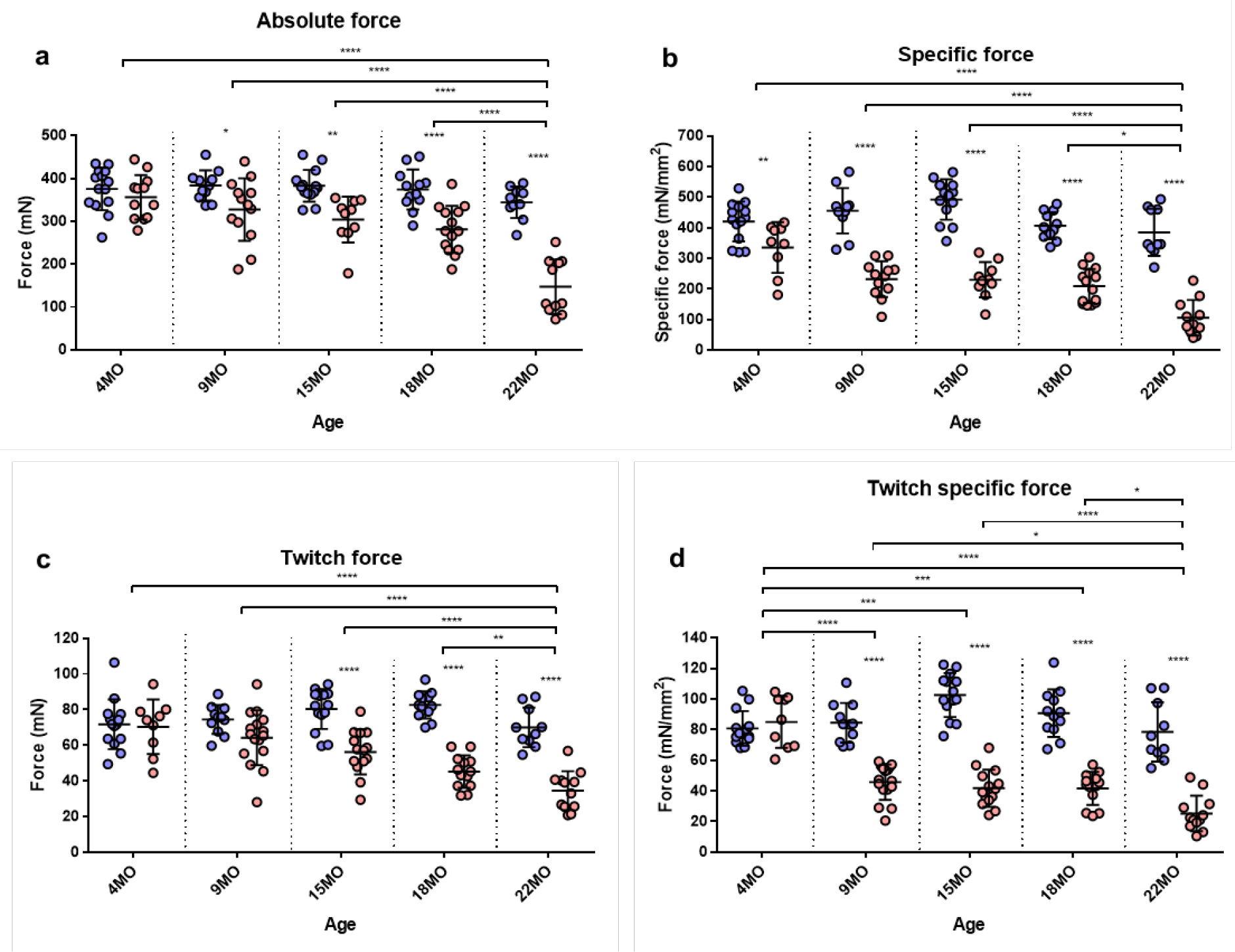
Maximum tetanic and twitch force. **a)** Scatterplot of the maximum absolute force generated by EDL muscles across 4 month (n=14 control; n=12 *mdx*), 9 month (n=11 control; n=13 *mdx*), 15 month (n=14 control; n=10 *mdx*), 18 month (n=12 control; n=14 *mdx*) and 22 month (n=10 control; n=11 *mdx*) age groups. **b)** Scatterplot of the maximum specific force (force per physiological cross sectional area) generated by EDL muscles across 4 month (n=14 control; n=9 *mdx*), 9 month (n=11 control; n=12 *mdx*), 15 month (n=14 control; n=10 *mdx*), 18 month (n=12 control; n=14 *mdx*) and 22 month (n=10 control; n=11 *mdx*) age groups. **c)** Scatterplot of the twitch force generated by EDL muscles across 4 month (n=14 control; n=9 *mdx*), 9 month (n=11 control; n=16 *mdx*), 15 month (n=14 control; n=14 *mdx*), 18 month (n=12 control; n=13 *mdx*) and 22 month (n=10 control; n=12 *mdx*) age groups. **d)** Scatterplot of the twitch specific force generated by EDL muscles across 4 month (n=14 control; n=9 *mdx*), 9 month (n=11 control; n=16 *mdx*), 15 month (n=14 control; n=14 *mdx*), 18 month (n=12 control; n=13 *mdx*) and 22 month (n=10 control; n=12 *mdx*) age groups. In each group the horizontal line indicates the mean value ± SD. Statistical differences between age groups are displayed above the graph, statistical differences displayed within graphs are differences between genotypes assessed by two-way ANOVA, *post hoc* analysis using Sidak’s multiple comparisons test. ****P<0.0001, ***0.0001<P<0.001, **0.001<P<0.01, *0.01<P<0.05.

Twitch forces for all groups are shown in Figure 3c. No significant differences in twitch force were seen between genotypes in both 4 month and 9 month old mice, however as the animals age the dystrophic muscles produce significantly less force than age matched littermate control muscles. Differences in EDL twitch force between aged *mdx* mice and littermate controls; 15 months ∼30% (MD -24.04, 95% CI [-35.8, -12.28], P<0.0001), 18 months ∼45% (MD -37.38, 95% CI [-49.83, -24.92], P<0.0001) and 22 months ∼51% (MD -35.51, 95% CI [-48.84, -22.19], P<0.0001). Again the 22 month old dystrophic cohort produced significantly less starting twitch force compared to all other age groups (F (4, 115)=4.99, P<0.0001). When twitch force values were corrected for cross sectional area in Figure 3d the 4 month dystrophic EDL produced the same specific force as littermate controls, however, despite being heavier the dystrophic EDL muscles produced less specific force from 9-22 months (F (4, 115)=20.8, P<0.0001), which again correlated with the increase in the number and complexity of branched fibers during this period Figure 1.

### Force frequency parameters are not significantly different with respect to age and genotype

Force frequency curves were generated from EDL muscle recordings at various frequencies and curve fitted using the sigmoidal equation given in the methods. These curves were generated from data at all age groups and genotypes (Figure 4a), however for clarity data at 4 months and 22 months old have been isolated in Figure 4b to highlight the shape and slope of the force frequency curves. There were no significant differences in half frequency and hill coefficient with respect to genotype or age (Table 1) consistent with previous studies. No differences in force frequency parameters are consistent with earlier reports of ^12, 25, 26^ no major fiber type changes with regard to myosin heavy chain isoforms or Ca^2+^ binding affinity of troponin resulting from the absence of dystrophin or age. Figure 4a provides a dramatic illustration of the decline in force with age in dystrophic *mdx* EDL and its alikeness to the development of branched fibers within the same cohorts (Figure 1).

**Figure 4:**
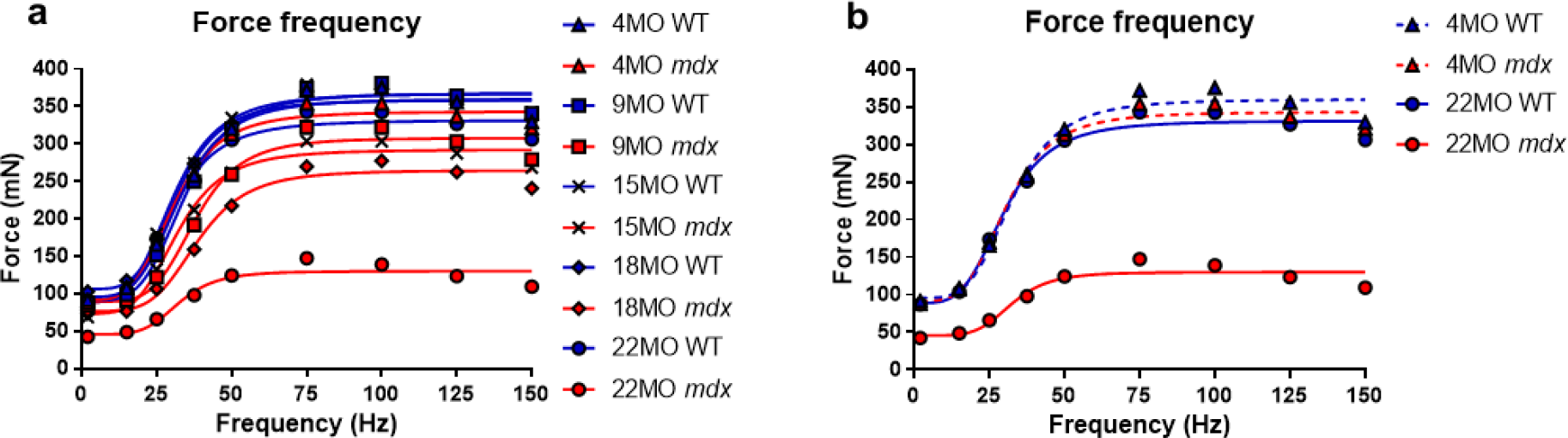
Force frequency curves. **a)** Aggregated force frequency curves from EDL muscles to visualize differences between genotype across 4 month (n=7 control; n=7 *mdx*), 9 month (n=6 control; n=7 *mdx*), 15 month (n=7 control; n=6 *mdx*), 18 month (n=6 control; n=7 *mdx*) and 22 month (n=6 control; n=7 *mdx*) age groups. Muscle absolute force was measured at different stimulation frequencies and a sigmoidal curve (lines) was fitted to the data points (see methods), contractile properties were generated from these curves (see table 1). **b)** For clarity 4 month and 22 month old age groups have been isolated to highlight the shape and slope of these force frequency curves. Data shown in these curves are mean only, SD was omitted for visual clarity.

### Increased isometric force loss in mdx with age

The absolute force of each of the series of 11 isometric contractions was plotted in figure 5a to visualize force decline throughout the isometric protocol and recovery after 5 minutes.

**Figure 5:**
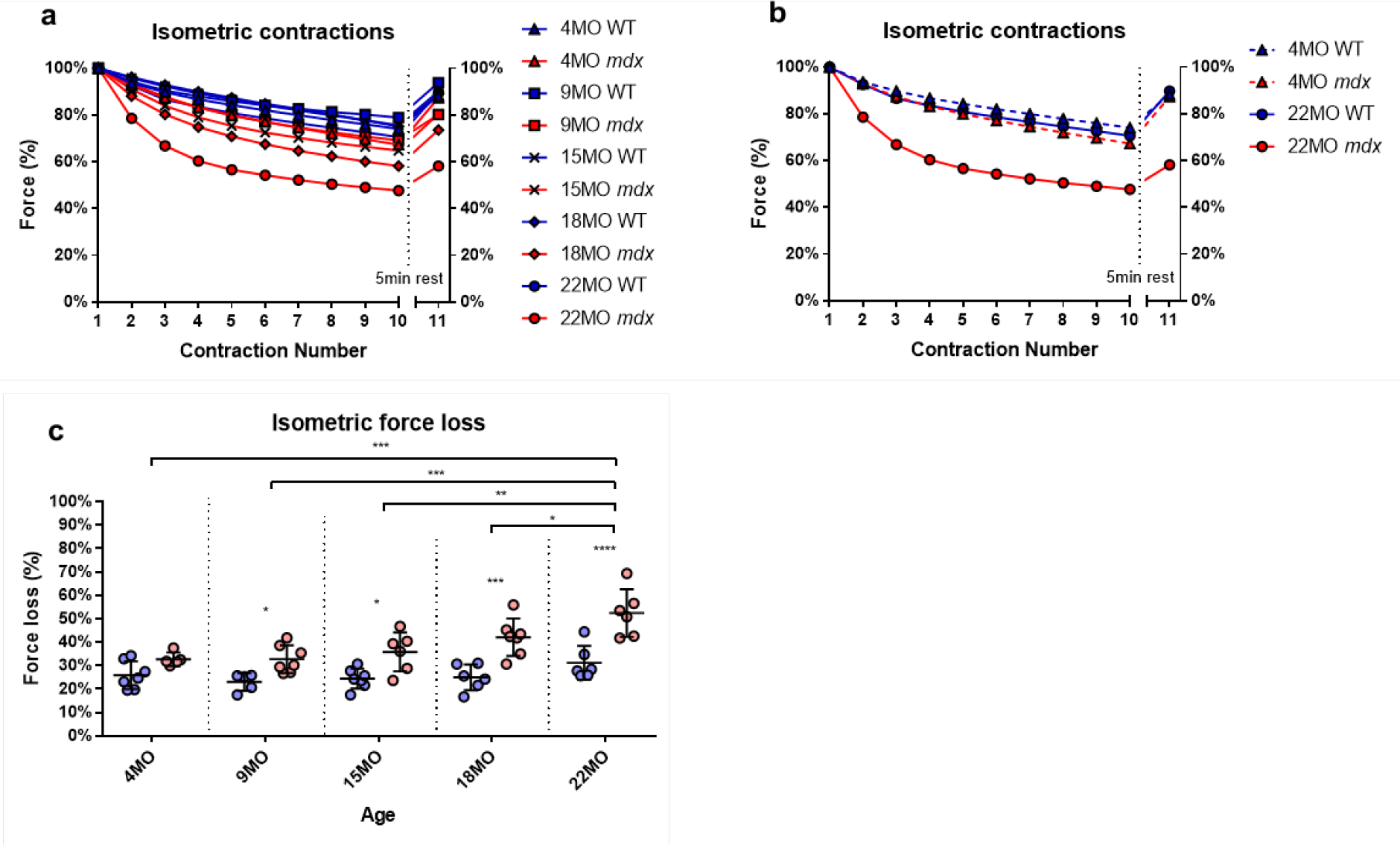
Percentage force changes throughout repeat isometric contractions and recovery. **a)** Shows the change in isometric force expressed as a percentage of starting force across 10 consecutive contractions and an 11^th^ recovery contraction after 5 minutes for all age groups. **b)** For clarity 4 month and 22 month old age groups have been isolated to highlight the differences in isometric force loss and recovery across these contractions. Data shown in these curves are mean only; SD was omitted for visual clarity. **c)** A scatterplot of the cumulative force loss by the EDL across 4 month (n=7 control; n=5 *mdx*), 9 month (n=5 control; n=7 *mdx*), 15 month (n=7 control; n=6 *mdx*), 18 month (n=6 control; n=7 *mdx*) and 22 month (n=6 control; n=6 *mdx*) age groups (Note: these numbers apply to a-c). In each scatterplot group the horizontal line indicates the mean value ± SD. Statistical differences between age groups are displayed above the graph, statistical differences displayed within graphs are differences between genotypes assessed by two-way ANOVA, *post hoc* analysis using Sidak’s multiple comparisons test. ****P<0.0001, ***0.0001<P<0.001, **0.001<P<0.01, *0.01<P<0.05.

Figure 5b isolates 4 month and 22 month age groups to highlight the upper and lower ends of force loss in dystrophic muscles compared to age matched controls. More isometric force is lost in dystrophic EDL muscles as the animals age (Figure 5c), dystrophic muscles from mice at 4 months lost similar amounts of force compared to littermate controls. The vulnerability of these *mdx* muscles to repeated isometric induced contractions progressively increased with age, greatest at the 22 month old age group where *mdx* muscles lost ∼52% of their starting force c.f. ∼31% in control animals (MD 21.28, 95% CI [11.09, 31.48], P<0.0001). The rest period between each isometric contraction is enough to ensure there is no significant fatigue, and indeed all control muscles recover close to 100% of their starting force after a 5 minute rest. The 22 month old *mdx* only recover to 60%, suggesting the isometric contractions have ruptured branched fibers (Figure 1). Once branching is present in the *mdx* EDL there is a correlation between the isometric force loss and the degree of fiber branching Figure 5a compared to Figure 1.

### Dystrophic EDL force loss per eccentric contraction was gradual for 4 month mice but sudden in all other age groups

The isometric force drops after each EC are expressed as a portion of starting force for all age groups in Figure 6a. Figure 6b omits traces for 9, 15 and 18 month old mice for clarity in distinguishing the gradual force drop seen in 4 months old *mdx* animals and sharp sudden drop in aged dystrophic animals. In all age groups, *mdx* EDL muscles lost more force during the EC protocol than age matched littermate control counterparts. However, the *mdx* EDL muscles in the 4 month group were the only dystrophic muscles to lose force gradually throughout the 6 ECs resembling the step-like declines seen in control muscles (Figure 6a), losing approximately ∼44% of starting force c.f. ∼18 in controls on the first contraction (MD 25.4, 95% CI [17.62, 33.18], P<0.0001). In comparison, all dystrophic EDL muscles in other age groups lost at ≥80% of their starting force after the first EC with the greatest lost at ∼92% for 22 month old *mdx* mice c.f. ∼16% in controls (MD 76.29, 95% CI [68.05, 84.54], P<0.0001) (Figure 6c).

**Figure 6:**
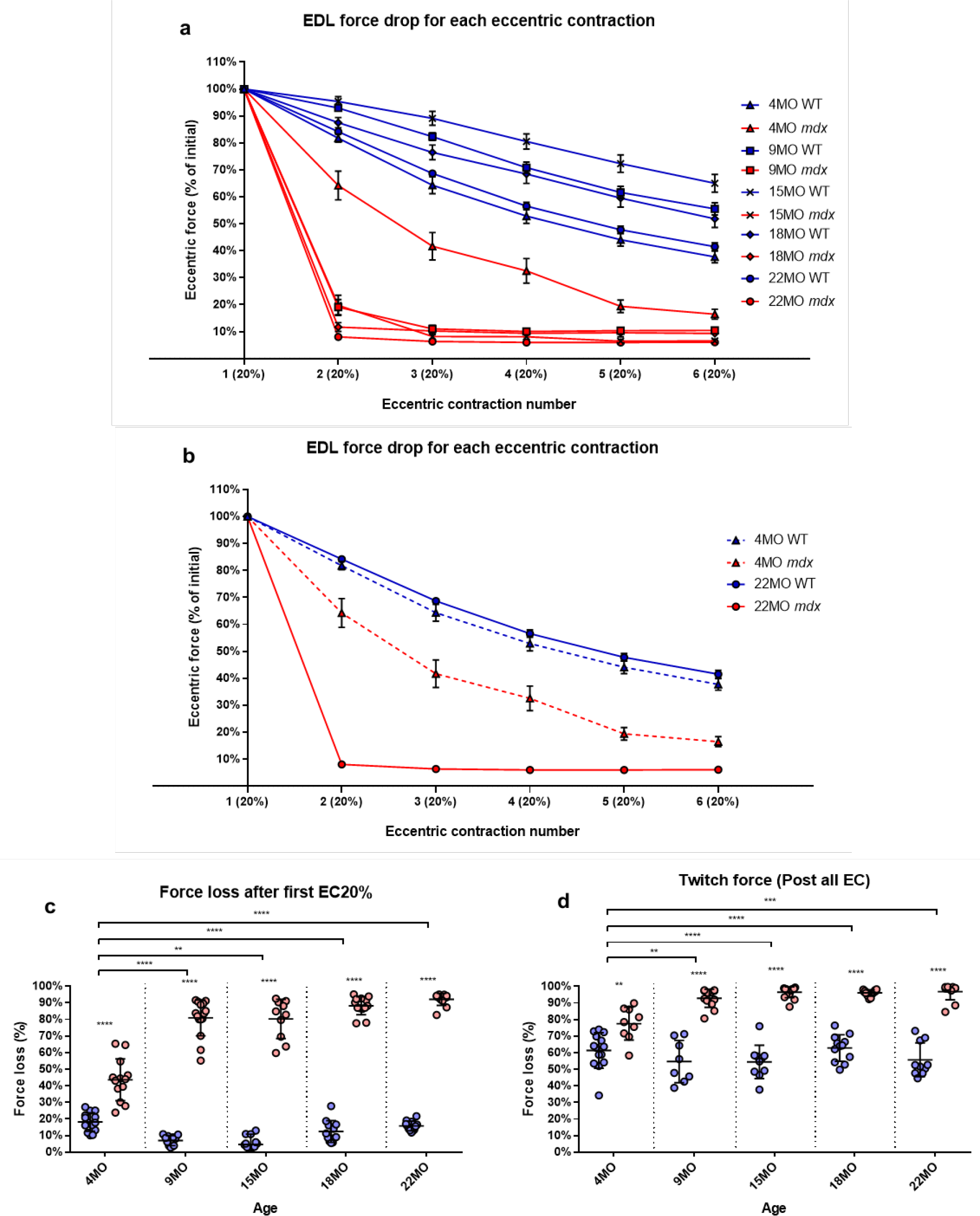
Percentage force loss resulting from a series of six ECs at 20% excursion from L_o_. Forces were normalized in each group with P_max_=100%. **a)** XY plot of EDL force loss for *mdx* mice and littermate controls across all age groups. **b)** For clarity 4 month and 22 month old age groups have been isolated to highlight the differences in eccentric force loss across these contractions. Data shown in these curves are mean ± SD. **c)**. Scatterplots of the force loss after the first 20% EC across 4 month (n=14 control; n=13 *mdx*), 9 month (n=11 control; n=15 *mdx*), 15 month (n=12 control; n=10 *mdx*), 18 month (n=12 control; n=13 *mdx*) and 22 month (n=12 control; n=12 *mdx*) age groups (Note: these numbers apply to **a-c**). **d)** Scatterplots of the twitch force loss after all six EC across 4 month (n=14 control; n=9 *mdx*), 9 month (n=11 control; n=16 *mdx*), 15 month (n=14 control; n=14 *mdx*), 18 month (n=12 control; n=13 *mdx*) and 22 month (n=10 control; n=12 *mdx*) age groups. For scatterplots in **c & d**, the horizontal line indicates the mean value ± SD. Statistical differences between age groups are displayed above the graph, statistical differences displayed within graphs are differences between genotypes assessed by two-way ANOVA, *post hoc* analysis using Sidak’s multiple comparisons test. ****P<0.0001, ***0.0001<P<0.001, **0.001<P<0.01, *0.01<P<0.05.

At the end of the 6 EC protocol; 4 month *mdx* muscles lost a total of ∼84% of initial force compared to ∼62% in control muscles (MD 21.25, 95% CI [14.2, 28.31], P<0.0001), this drop in control muscle force was previously reported in young 6-9 week control muscles in Kiriaev, et al. ^12^. Whereas 15 month *mdx* muscles lost ∼94% of initial force compared to ∼35% in control muscles (MD 58.49, 95% CI [50.65, 66.33], P<0.0001) similar to 22 month *mdx* muscles that lost ∼94% of initial force compared to ∼59% in control muscles (MD 35.81, 95% CI [28.38, 43.29], P<0.0001) (Figure 6a).

In line with the tetanic force data the 9-22 month dystrophic EDL twitch force twitch force dropped dramatically after the entire EC protocol (Figure 6d). EDL muscles from 4 month *mdx* animals lost an average of ∼78% of their starting twitch force c.f. ∼61% in controls after the EC protocol (MD 16.19, 95% CI [6.92, 25.45], P=0.0016), 9 month *mdx* animals lost ∼93% of their starting twitch force c.f. ∼55% in controls after the EC protocol (MD 38.16, 95% CI [28.48, 47.83], P<0.0001). In the aged *mdx* cohorts, EDL muscle twitches barely produced any force reflecting the consequence to force loss due to EC induced damage in Figures 6a-c. Again the largest force deficit occurred in 22 month *mdx* mice losing ∼97% of the pre-EC twitch force c.f. ∼56% in littermate controls (MD 41.14, 95% CI [32.06, 50.21], P<0.0001). Once again, the correlation between the catastrophic force loss experienced by the 9-22 month *mdx* cohort and increases in fiber branching is striking.

### Rapid recovery from eccentric force loss is seen in 4 month mdx but to a lesser degree as the animal ages

Using the rational of Olthoff, et al. ^44^ we gave the post EC muscles 2 hours rest to see the amount of recovery attributable to non-sarcolemmal damaged muscle fibers (Figure 7a). Our EC was strong enough to cause fiber damage in control mice where there was a 20-35% non-recoverable force loss. Figure 7b illustrates the EDL recovery post EC at 20 minutes intervals for 4 month and 22 month old EDL muscles. In the *mdx* cohort, 4 month old EDL muscles showed the greatest amount of recovery of up to ∼47% of their starting force c.f. ∼68% in age matched littermate controls (MD -22.01, 95% CI [-34.36, -9.66], P<0.0001) whereas the 22 month old *mdx* EDL muscles recovered the least ∼25% of starting c.f. ∼65% in age matched littermate controls (MD -40.29, 95% CI [-52.88, -27.69], P<0.0001). The recovery in *mdx* EDL decreases as the dystrophic animal ages resulting in a decline in end recovered force as shown in Figure 7c. Dystrophic EDL muscles at 9 months recovered ∼41% (MD -38.63, 95% CI [-51.06, -26.2], P<0.0001), 15 months recovered ∼29% (MD -44.45, 95% CI [-59.08, - 29.82], P<0.0001) and 18 months recovered ∼33% (MD -39.08, 95% CI [-51.43, -26.73], P<0.0001) of starting force. Given both 4 month *mdx* mice and 22 month *mdx* mice contain 100% regenerated dystrophin-negative muscle fibers ^13^. We attribute the large non-recoverable force loss in old *mdx* EDL to the increase in fiber branching (Figure 1).

**Figure 7:**
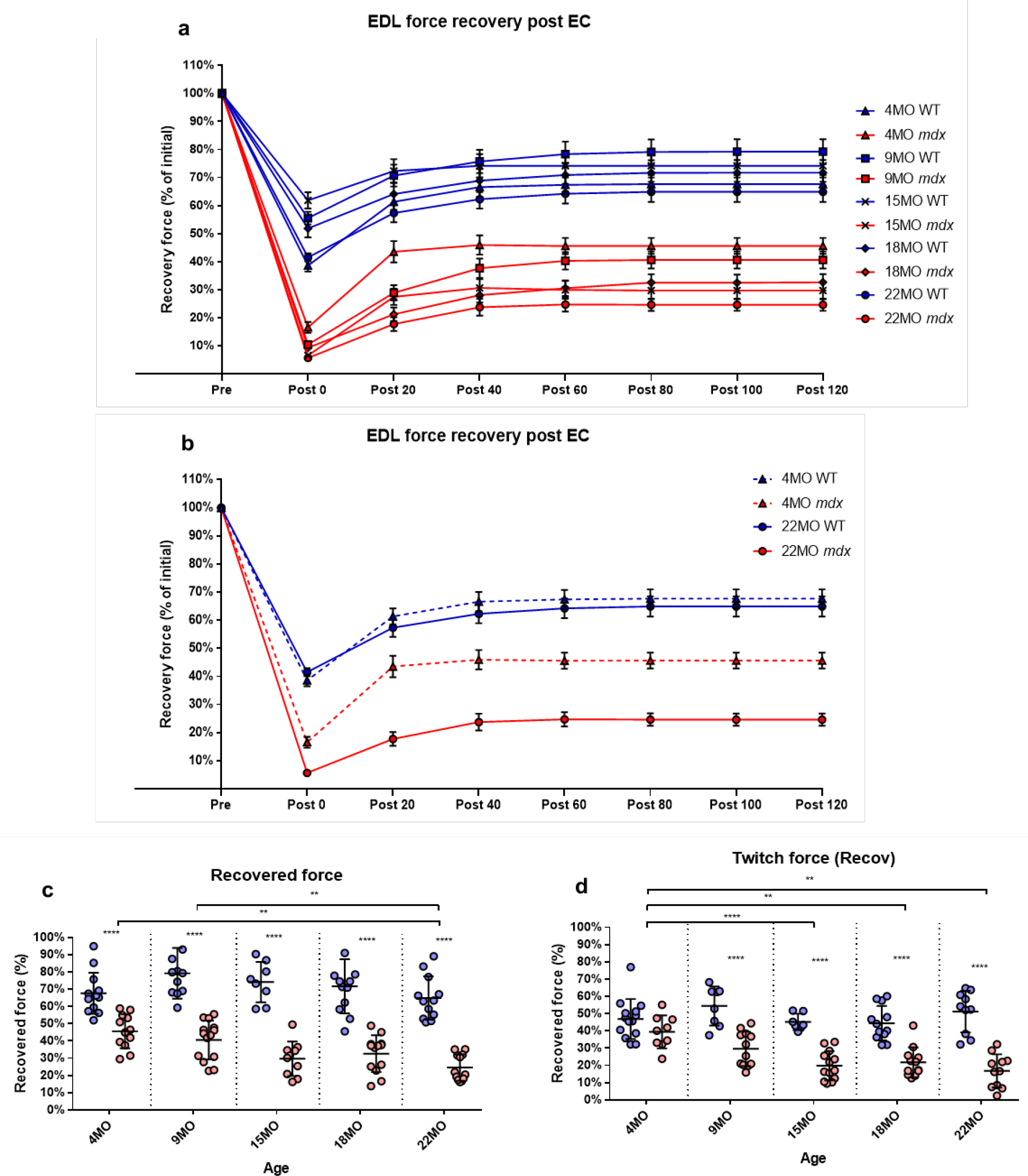
Percentage force recovery for up to 120 minutes post EC protocol. Forces were normalized in each group with P_max_=100%. **a)** XY plot of post EC EDL force recovery over 120 minutes at 20 minute intervals for *mdx* and littermate controls across all age groups. **b)** For clarity 4 month and 22 month old age groups have been isolated to highlight the differences in post EC recovery across these contractions. Data shown in these curves are mean ± SD. **c)** Scatterplots of the recovery at 120 minutes across 4 month (n=13 control; n=12 *mdx*), 9 month (n=11 control; n=14 *mdx*), 15 month (n=8 control; n=10 *mdx*), 18 month (n=12 control; n=13 *mdx*) and 22 month (n=12 control; n=12 *mdx*) age groups (Note: these numbers apply to **a-c**). **d)** Scatterplots of the twitch force recovery at 120 minutes across 4 month (n=14 control; n=9 *mdx*), 9 month (n=11 control; n=12 *mdx*), 15 month (n=7 control; n=13 *mdx*), 18 month (n=12 control; n=12 *mdx*) and 22 month (n=10 control; n=11 *mdx*) age groups. For scatterplots in **c & d**, the horizontal line indicates the mean value ± SD. Statistical differences between age groups are displayed above the graph, statistical differences displayed within graphs are differences between genotypes assessed by two-way ANOVA, *post hoc* analysis using Sidak’s multiple comparisons test. ****P<0.0001, **0.001<P<0.01.

In regards to twitch force recovery (Figure 7d), at 4 months of age there were no significant differences between dystrophic and control muscles. This is a key finding, along with the fact that at 4 months there is no difference in either absolute or specific twitch force (Figure 3c,d), and the correlation with the low number and reduced complexity of branching at this age (Figure 1). However, the ability for EDL muscles from dystrophic animals to recover twitch force after our EC protocol decreased over the 9-22 month old age groups (Figure 7d). EDL twitch force recovery for adult and aged groups; 9 months ∼30% for *mdx* c.f. ∼54% for controls (MD -23.86, 95% CI [-36.7, -13.03], P<0.0001), 15 months ∼20% for *mdx* c.f. ∼45% for controls (MD -25.34, 95% CI [-37.49, -13.18], P<0.0001), 18 months ∼22% for *mdx* c.f. ∼44% for controls (MD -22.66% CI [-33.24, -12.07], P<0.0001) and 22 months ∼17% for *mdx* c.f. ∼51% for controls (MD -34.35, 95% CI [-23.02, -45.69], P<0.0001). Once again it is striking that although 4-22 month old dystrophic EDL muscles contain regenerated dystrophin negative fibers, that the force loss and EC deficit is strongly correlated with the degree and complexity of branched fibers (Figure 1).

### Stiffness

The EDL muscle stiffness was calculated during the first EC and reported in Figure 8a. In the 4 month old group, there was no significant difference in stiffness between *mdx* EDL and age matched littermate control EDL muscles, in contrast age groups 9-22 months show a significant increase in stiffness compared to age matched littermate control animals. The increase in stiffness is correlated with the increase in branching (Figure 1) and is likely the consequence of branching effectively increasing the number of small muscle fibers arranged in series within the aging dystrophic muscle. Differences in muscle stiffness between *mdx* and controls for each of these age groups are; 9 months (MD 74.38, 95% CI [55.55, 93.21], P<0.0001), 15 months (MD 80.97, 95% CI [60.66, 101.3], P<0.0001), 18 months (MD 57.48, 95% CI [37.17, 77.78], P<0.0001) and 22 months (MD 29.11, 95% CI [9.75, 48.47], P=0.0007).

**Figure 8:**
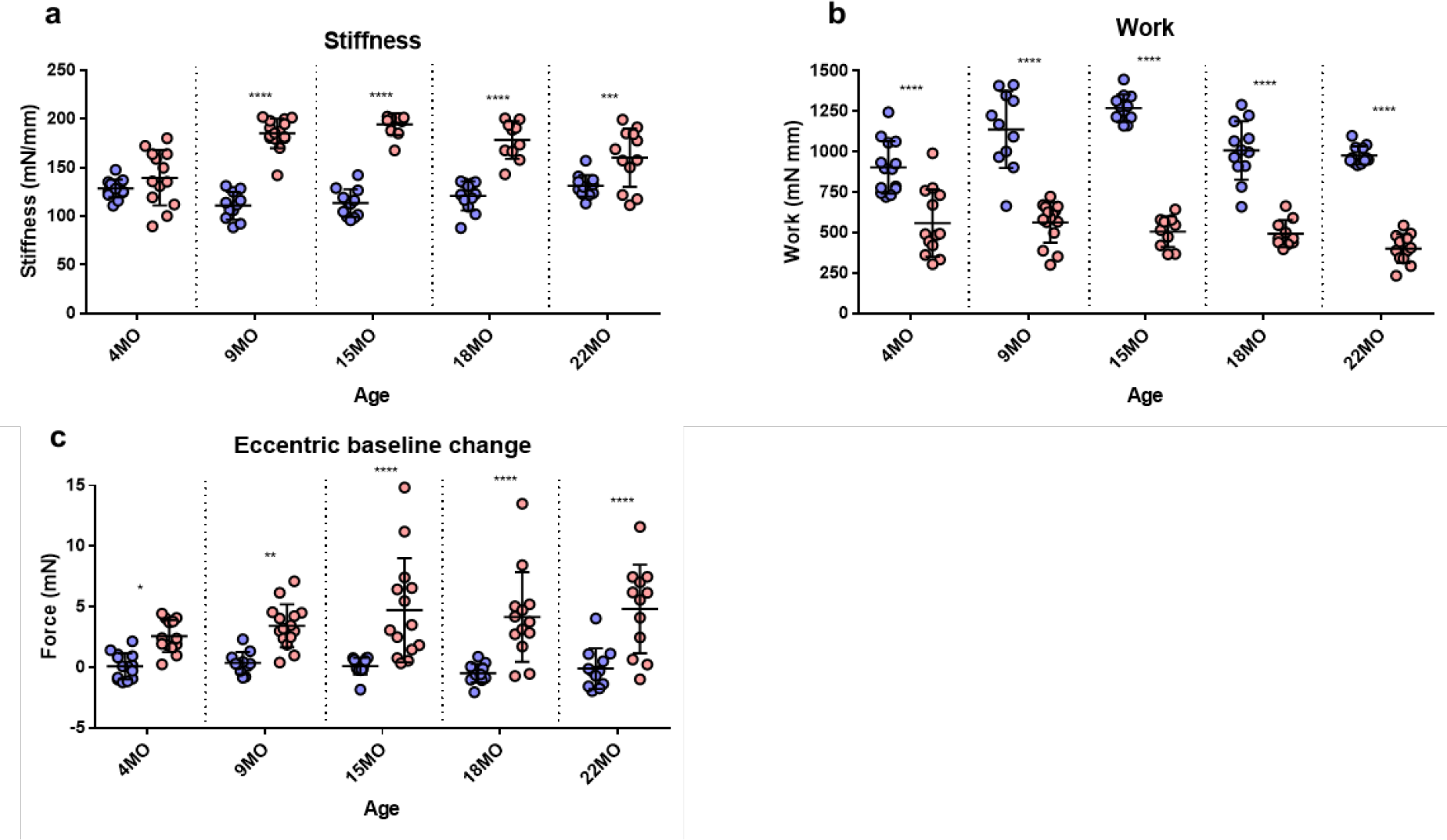
Stiffness, Work done and baseline force. **a)** Scatterplots showing the EDL muscle stiffness calculated during the first 20% EC across 4 month (n=14 control; n=13 *mdx*), 9 month (n=11 control; n=15 *mdx*), 15 month (n=12 control; n=10 *mdx*), 18 month (n=12 control; n=10 *mdx*) and 22 month (n=12 control; n=12 *mdx*) age groups. **b)** Scatterplots showing the work done by EDL muscles calculated during the first 20% EC across 4 month (n=14 control; n=13 *mdx*), 9 month (n=11 control; n=15 *mdx*), 15 month (n=12 control; n=10 *mdx*), 18 month (n=12 control; n=10 *mdx*) and 22 month (n=12 control; n=12 *mdx*) age groups. **c)** Scatterplots showing the baseline force change post EC for *mdx* and littermate control EDL muscles across 4 month (n=14 control; n=13 *mdx*), 9 month (n=11 control; n=14 *mdx*), 15 month (n=9 control; n=14 *mdx*), 18 month (n=12 control; n=10 *mdx*) and 22 month (n=12 control; n=12 *mdx*) age groups. In each scatterplot group the horizontal line indicates the mean value ± SD. Statistical differences between age groups are displayed above the graph, statistical differences displayed within graphs are differences between genotypes assessed by two-way ANOVA, *post hoc* analysis using Sidak’s multiple comparisons test. ****P<0.0001, ***0.0001<P<0.001, **0.001<P<0.01, *0.01<P<0.05.

### Work done

We calculated the work done as the area under the force curve during the active lengthening phase of the EC (Figure 8b) and illustrates that the work done during the EC was significantly greater in age matched littermate controls compared with the dystrophic EDL muscle. Differences in work done between *mdx* and controls for each age group are; 4 months (MD - 334, 95% CI [-490.2, -197.8], P<0.0001), 9 months (MD -574.9, 95% CI [-725.6, -424.2], P<0.0001), 15 months (MD -763.5, 95% CI [-926, -600.9], P<0.0001), 18 months (MD - 514.1, 95% CI [-676.6, -351.6], P<0.0001) and 22 months (MD -576.1, 95% CI [-731, - 421.1], P<0.0001).

### Increase in resting force after the first EC

The change in resting baseline absolute force after the first EC is shown in Figure 8c. In littermate control EDL muscles, baseline force remained at zero (the set point prior to the EC protocol) throughout different age groups following the first EC. In *mdx* EDL muscles after an EC produced a significantly higher resting force across all age groups, which increased in magnitude and spread with age. Differences in baseline change between *mdx* and controls for each age group are; 4 months (MD 2.482, 95% CI [0.09, 4.88], P=0.039), 9 months (MD 3.06, 95% CI [0.60, 5.53], P=0.0077), 15 months (MD 4.61, 95% CI [2.23, 6.96], P<0.0001), 18 months (MD 4.64, 95% CI [2.15, 7.13], P<0.0001) and 22 months (MD 4.93, 95% CI [2.39, 7.47], P<0.0001).

### Twitch

As pointed out by Peczkowski, et al. ^81^ while maximal isometric tetanic contractions are most commonly used to assess and report muscle function *in vitro*, *in vivo* muscles likely contract at sub-maximal levels. In order to address this, we looked at twitch kinetics as an outcome parameter. Twitch kinetics were measured (1) pre-eccentric, (2) post-eccentric and (3) after recovery from the EC protocol with statistical analysis presented in Table 1.

### Twitch half relaxation times

Twitch half relaxation times did not significantly change in littermate controls throughout all three measures across all age groups, (1) pre-eccentric, this was the same for the dystrophic group with the exception of 4 month old *mdx* EDL which took significantly longer to relax. Following ECs (2) post-eccentric twitch relaxation time for *mdx* EDL muscles increased in all age groups and were significantly higher than control values. When measured (3) after recovery from the EC protocol, dystrophic EDL muscles had reduced half relaxation time closer to starting values but still took significantly longer to relax relative to age matched littermate controls (see Table 1 for genotype statistics and distribution for kinetics).

### Twitch time to peak

Except for 4 month old dystrophic EDLs (1) pre-eccentric twitch time to peak measures showed that dystrophic EDL muscles had a significantly faster twitch time to peak compared to age matched controls in all age groups. In the 4 month old group, both *mdx* and control muscle reached peak contraction at a similar time. Most likely due to a consequence in the different levels of absolute force, see Figure 3c, (if the muscle produces a greater twitch force it will take a longer time to achieve this if other parameters are the same). (2) post-eccentric 4 month old *mdx* EDL muscles TTP was significantly slower time than controls. The time to peak for all older age groups remained similar with no differences between age or genotype. (3) After recovery from the EC protocol dystrophic EDL muscles showed a significantly slower TTP than control animals at 4, 9 and 22 month age groups and remained similar for remaining cohorts (Table 1), this is interesting because given the lower forces generated by the dystrophic muscles c.f. age matched littermate controls (Figure 7d) at 9-22 months we would predict dystrophic animals should have the faster TTP while by the same rational 4 month groups should be the same TTP.

### RT50/TTP ratio for twitch kinetics

Peczkowski, et al. ^81^ developed the RT50/TTP ratio as a way of assessing twitch kinetics between genotypes and ages. Using this measure, littermate control mouse EDL muscles remain unchanged due to EC induced injury/recovery and stayed consistent across all control age groups (Table 1) and treatments (1) pre-eccentric, (2) post-eccentric and (3) after recovery from the EC protocol (with the EDL muscles contracting at a slower rate than they relax (RT50/TTP ratio less than 1). (1) Pre-eccentric RT50/TTP ratio was similar for dystrophic muscles c.f. littermate controls in most age groups (Table 1). (2) Post-eccentric RT50/TTP ratio increased for all *mdx* EDL muscles across all groups and remained elevated when measured (3) after recovery from the EC protocol.

The similar RT50/TTP ratio (1) pre-eccentric in *mdx* and age matched littermate controls suggests the fiber type profiles of the EDL remains unchanged in the *mdx*, while the higher RT50/TTP ratio reported for dystrophic muscle fibers post EC shows dystrophic muscles contracting at a similar rate to relaxing possibly due to the presence of stressed branched fibers with reduced excitability and slower rates of contraction and relaxation throughout the branched syncytium.

### Light microscope morphology of enzymatically isolated single fibers

Representative stitched images of intact muscle fibers taken at magnification (X100) on a light microscope demonstrating various degrees of complex fiber branching in the senescent (22 month) *mdx* EDL (Figure 9). Fiber a) shows an example of a simple branched fiber containing one branched end and multiple splits within itself that develop along the length of the fiber. These splits along the fiber can become quite large and more noticeable such as in Fiber b) where we can see several branched offshoots from the main fiber trunk. Fibers c) & d) shows examples of complex branching (4+ branches) with multiple offshoots along the length of the dystrophic fiber. Branch points have been marked with arrows for each fiber.

**Figure 9:**
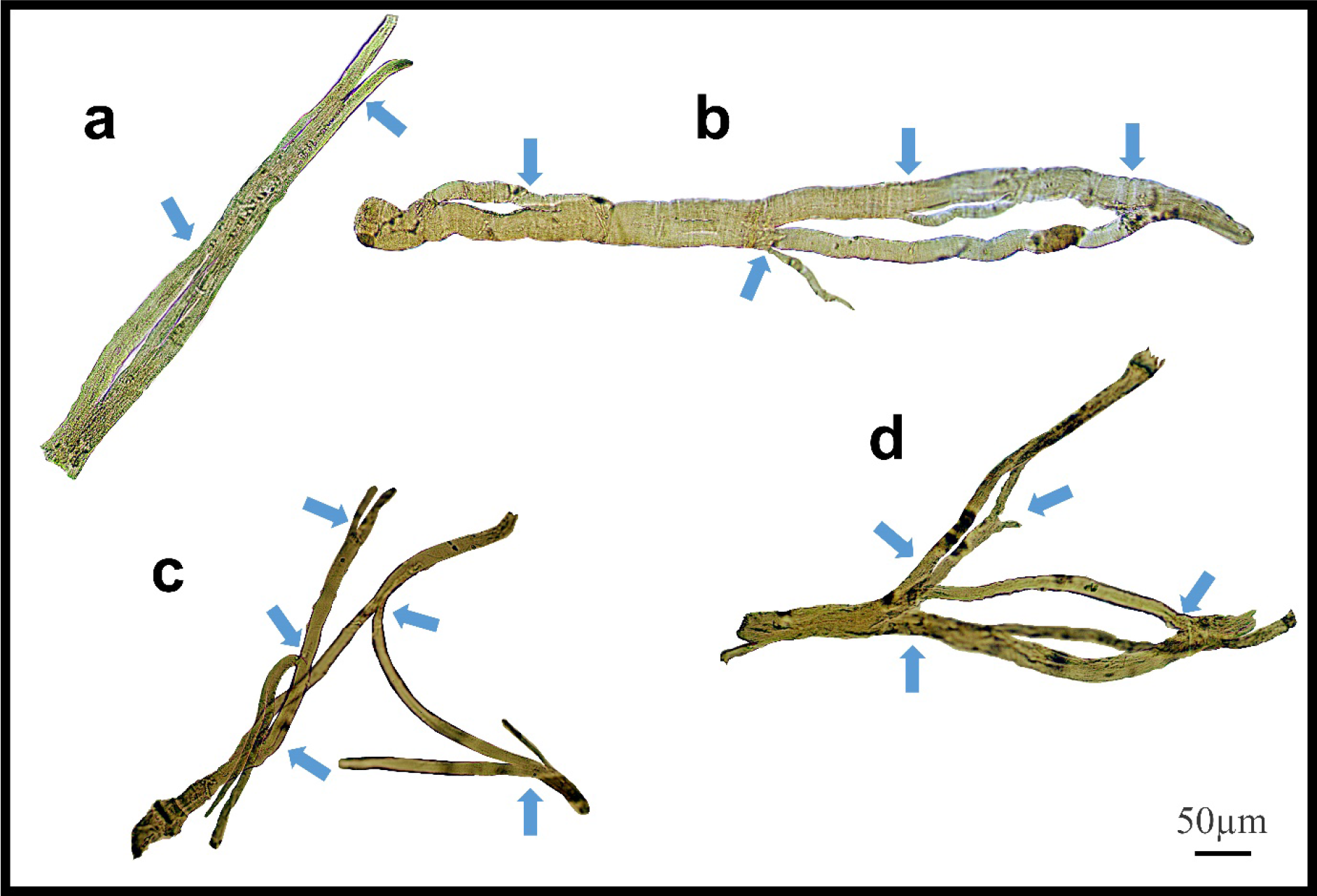
Examples of medium power light microscope images (X100) of enzymatically digested EDL muscle fibers taken from a single 22 month old dystrophic mouse. Note the fibers have been stitched together at the same magnification from photomicrographs taken from overlapping fields of view to capture a large portion of the fiber, scale bar provided at 50µm. Backgrounds debris from the digest process have been cleared to focus on the muscle fiber and arrows mark areas where fiber branching has occurred (To see examples where debris have not been removed see Kiriaev, et al. ^12^. **a)** A fiber containing a single branch with slits forming within the trunk. **b)** A fiber showing rejoining of these slits within the trunk and multiple offshoots off the edge of the muscle. **c & d)** Examples of complex branched muscle fibers.

Figure 10 provides some examples of (single field X100) light microscope images taken of branch points in aged *mdx* EDL muscle fibers before EC. The branch pattern in these *mdx* fibers range from offshoots from the main trunk of the fiber shown in panels a), c), g), i) & j), to splits that rejoin mid fiber in panels d) & e) as well as branching towards the end of fibers in panels b), f), h) & k).

**Figure 10:**
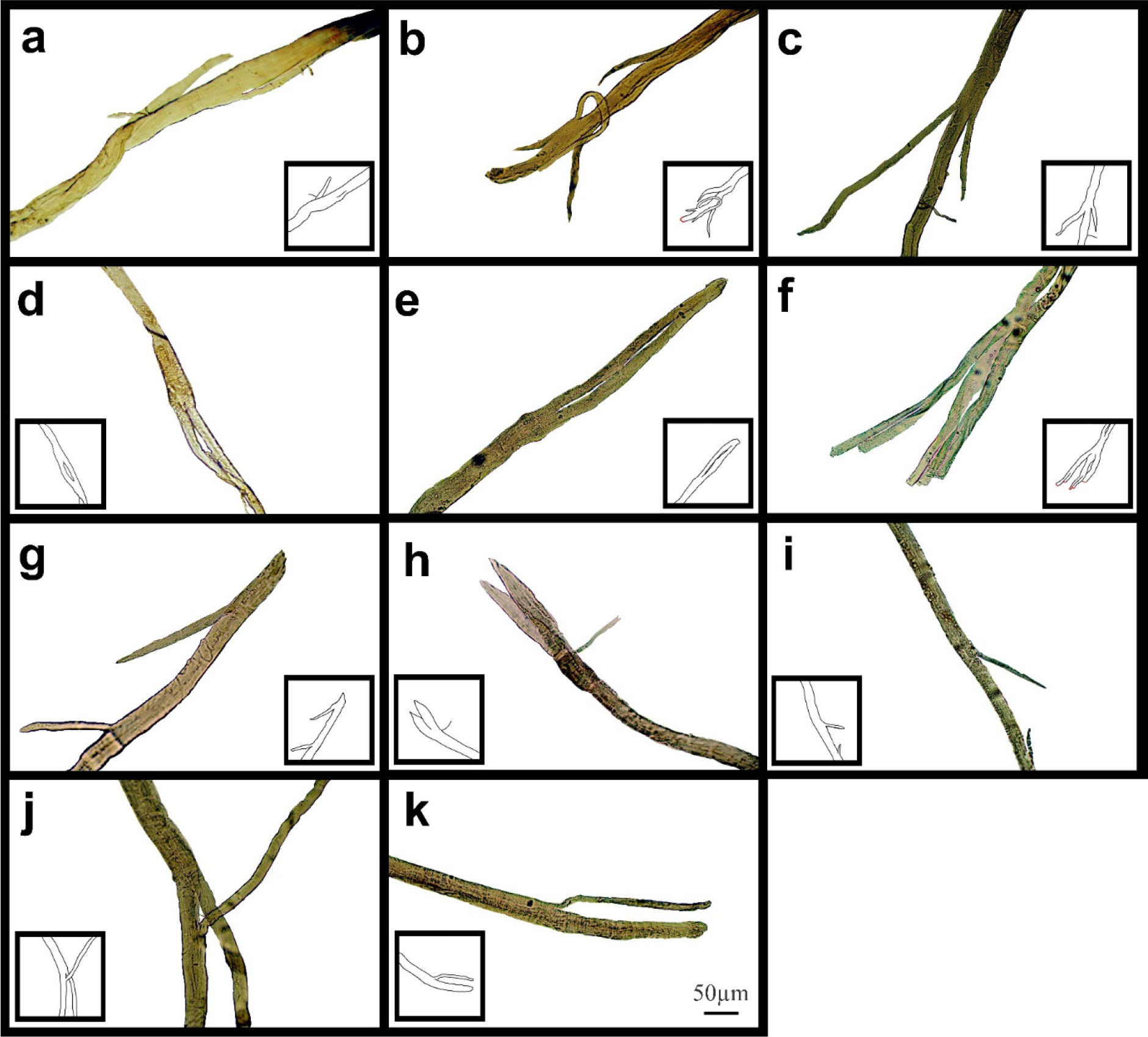
Examples of medium power light microscope images (X100) of muscle fiber branch point sections that have not undergone EC from adult and senescent *mdx* mice. Cartoon inserts have been added to help visualize the various examples of branching in each image. **a-f)** Are portions of fibers from a 22 month *mdx* mouse EDL. **g-k)** Are portions of fibers from a 18 month *mdx* mouse EDL. Scale bar provided at 50µm in **k)** applies to all of figure 10.

Figure 11 shows images of post EC broken EDL fibers from aged *mdx* mice taken at magnification (X100). Cartoon inserts of been added to each image to help identify broken fibers (Red) and necrotic areas (Yellow).

**Figure 11:**
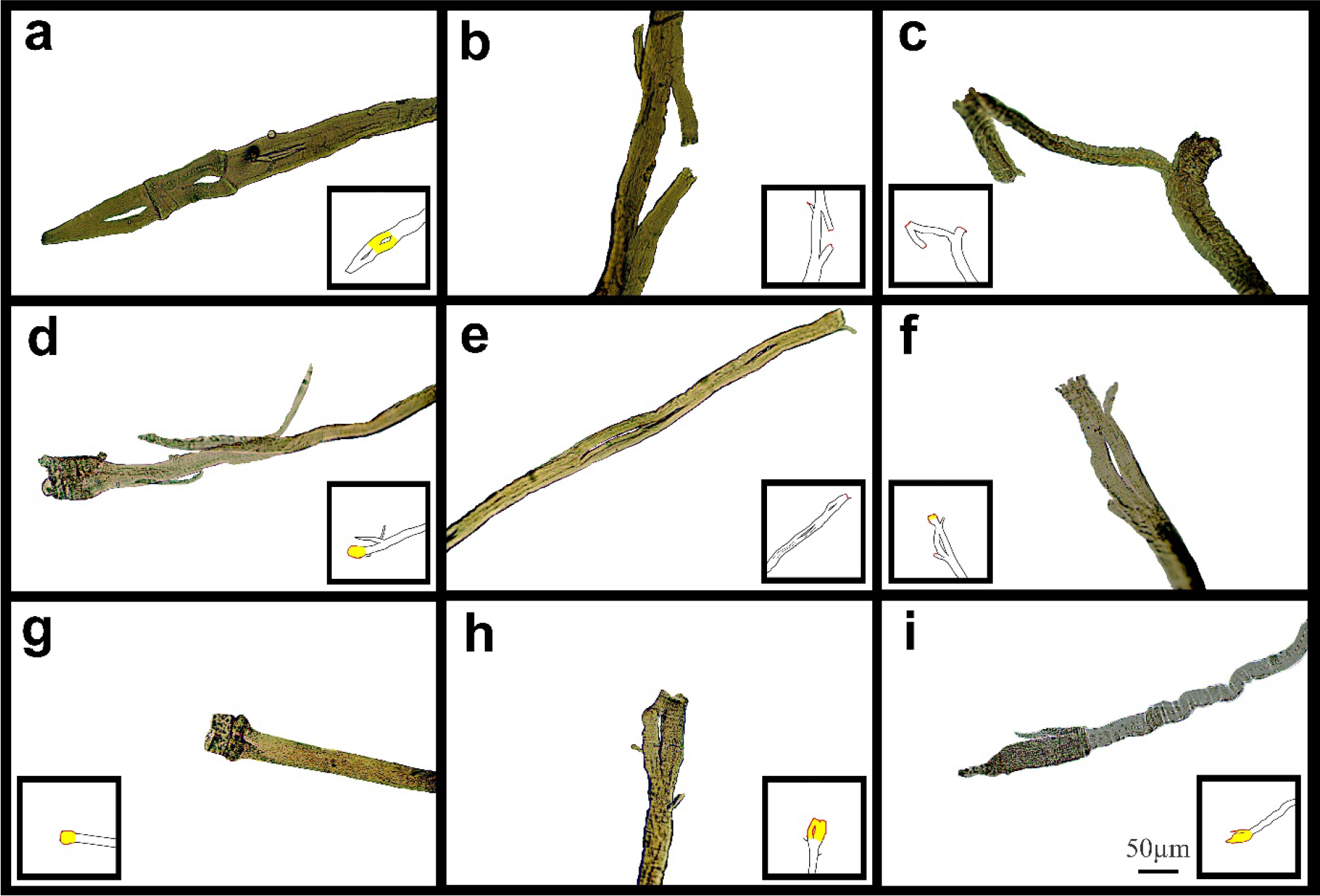
Examples of medium power light microscope images (X100) of broken muscle fiber sections that have undergone ECs from adult and senescent *mdx* mice. Broken areas have been outlined in red whilst swollen and necrotic areas highlighted yellow in cartoon inserts for each image. **a-d)** Are portions of fibers from a 22 month *mdx* mouse EDL. e-i) Are portions of fibers from a 18 month *mdx* mouse EDL. Scale bar provided at 50µm in **i)** applies to all of figure 11.

## DISCUSSION

The fast twitch EDL skeletal muscle *from the mdx* mouse, is the most used muscle to study the pathophysiology caused by the absence of dystrophin. The mouse EDL is a mix of fast fiber types; ∼79% type 2B (fast glycolytic), ∼16% type 2X and ∼4% type 2A (fast oxidative glycolytic)^82^. We have previously proposed a two-step model to describe the skeletal muscle pathology in the dystrophinopathies ^12, 15–17, 25^. Step-one; involves the absence of dystrophin triggering skeletal muscle fiber necrosis driven by a pathological increase in [Ca^2+^]_in_ likely caused by a combination of increased free radical damage and abnormal ion channel functioning. Step-two; involves repeated cycles of muscle fiber regeneration in response to step one, the regenerated dystrophin-deficient muscle fibers are abnormally branched, and this branching pathology increases in complexity with age as the number of regenerative cycles in step-2 increases. In this two-step model it is important to note that, step-one, will continue to occur at the same time as step-2, repeatedly triggering the cyclic nature of step-2. In older dystrophic animals we and others have proposed that it is the branched fast-twitch fibers, in and of themselves, which are the cause of the large irreversible EC force deficits in EDL muscles. Here we further test this two-stage model by looking at the correlation between fiber branching and the contractile pathology in dystrophin-deficient EDL muscles from *mdx* mice between 4 months to 22 months of age compared with age matched littermate controls. By 4 months of age 100% of *mdx* skeletal muscle fibers have undergone at least one round of necrosis/regeneration ^13^, so all the contractile findings reported here are from regenerated dystrophin deficient fast-twitch fibers. With reference to DMD our 4 and 9 month mice can be considered representative of the adolescent population while 15, 18 and 22 months represent adults and aged.

### Fiber branching

4 month to 22 month dystrophic EDL muscles are comprised of 100% regenerated dystrophin-negative muscle fibers ^6, 13, 55, 71, 74, 83, 84^, the major morphological change that occurs during this period is the formation of branched fibers which increase in number and complexity as the animal ages (Figures 1, 9)^8, 12, 13, 15, 25, 52, 54, 55, 74^. All the contractile deficits and increased dystrophic muscle weight we report in this study are correlated with the increase and complexity of these branches. In humans it has been proposed that skeletal muscle fiber branching is a key pathological sign of the progressive muscular dystrophies ^55, 85^. Branched skeletal fibers are found in the muscles of boys with DMD ^72, 73, 86–90^. In DMD boys extensive fiber branching has been correlated with a reduction in mobility ^72^. Since our first report ^8^ of fiber branching in the *mdx* limb muscles this finding has been confirmed by our laboratory and others^12,13,15,25,52-56,74^.

### Hypertrophy and loss of force in fast-twitch mdx muscles

Skeletal muscle hypertrophy is a characteristic of the dystrophinopathies ^17, 72, 84, 91–96^. In the mouse *mdx* EDL we and others have reported a 20-30% hypertrophy in EDL ^6, 12, 13, 18– 20, 24, 25, 49, 74, 97^ while reports of 20-60% hypertrophy have been published regarding the predominantly fast-twitch *mdx* TA ^10, 49, 71, 78, 98^. In the *mdx* mouse this hypertrophy initially maintains the absolute force output of the *mdx* EDL at 4 months, despite the larger muscles, the absolute force drops relative to age matched littermate control muscles (Figure 3a,c). In terms of maximum specific force the *mdx* EDL produces less at all ages (Figure 3b) compared to littermate controls and this difference increases with age, being most marked in the 22 month *mdx* EDL muscles. The picture is similar for the specific twitch force apart from the 4 month group where there is no difference (Figure 3d). Given the low turnover (step-2) of fibers at these ages and the strong correlation with the increase in number and complexity of branched fibers, we propose that the branching is responsible for both the hypertrophy and lower force output. The correlation between branching and force loss is also maintained when we look at the fitted force frequency curves (Figure 4) where the drop of force in *mdx* fibers is correlated with the increase in branched fibers (Figures 1, 9, 10).

Statistically there is no change in the half frequency or the Hill coefficient of the curves (Table 1) suggesting that functionally the *mdx* EDL muscles have remained fast-twitch during regeneration (4-22 months), this confirms studies that have reported no significant changes in fiber types with age in *mdx* muscle ^12, 25, 26, 69, 83, 99^. In the *mdx* limb muscles fiber branching itself has been shown to be the major component of muscle hypertrophy which occurs as the *mdx* mouse ages ^13, 55, 74^. Faber, et al. ^74^ demonstrates that in dystrophic EDL muscles a 28% increase in the number of fibers counted in transverse sections of muscles correlated with a 31% increase in myofiber branching. Furthermore, the study highlights the largest increase in myofiber number and branching both occurred at 12 weeks onwards confirming histologically a phase of severe degeneration occurring just prior. Recently a lifetime analysis of *mdx* skeletal muscle performed by Massopust, et al. ^49^ confirmed the findings of Faber, et al. ^74^ that myofiber diameter and volume develop normally until 12 weeks of age where dystrophic muscles become significantly larger than age matched controls. Importantly the study identified muscle fiber branching contributes to volume, diameter and CSA variability and branches themselves do not feature synapses thus receiving innervation from the original fiber^13,49,74^.

### Why does fiber branching reduce force output in mdx muscles?

Friedrich’s laboratory ^53^ have produced a convincing structural hypothesis where they propose the repeated bouts of regeneration and degeneration in *mdx* muscles produce branched fibers which have been structurally remodeled in such a way that contractile proteins have become misaligned and deviate from the long axis of the muscle, leading to asynchronized contraction and loss of force production. Furthermore, within a regenerated dystrophic fiber the group used second harmonic generation imaging to identify an increased variation in the angle of myofibril orientations ^56^ which they calculate could contribute to ∼50% of the progressive force loss with age in *mdx*. Additionally, several studies have confirmed that each branched fiber syncytium, no matter how many branches they contain (the most complex in excess of 10) is controlled by a single neuromuscular junction (a single motor nerve) ^49, 74, 100^. We have modelled ^16^ the transmission of a muscle action potential through a syncytium showing it will be slowed in small diameter branches and can fail to transmit when propagating is from a small diameter branch towards a large diameter branch, this means that the force output per cross-section of branched fiber syncytium will be reduced by the number of non-contracting small branches. Thus we propose the correlation of increased fiber branching with hypertrophy and decreased force output in dystrophin deficient muscles is not the result of the absence of dystrophin. Rather it is due to the absence of dystrophin triggering rounds of necrosis/regeneration which then produces branched fibers, that lead to muscle hypertrophy and reduce force output. Further evidence in support of this comes from our skinned fiber studies ^16, 51^ where we chemically remove the cell membrane (sarcolemma), thus removing the dystrophin protein link in control muscles and examine the force output from non-branched segments of contractile proteins from *mdx* EDL c.f. controls. We showed that there is no difference in the force output produced by these contractile proteins.

### 18 and 22 month dystrophic EDL muscle show an irreversible force deficit due to repeated maximal isometric contractions correlated with the number and complexity of branched fibers

When we gave the muscles 10 maximal isometric contractions, separated by rest periods of 60 seconds to reduce the impact of fatigue, it was only the 18 and 22 month old *mdx* muscles which showed a significant loss of force that did not recover. Our argument is summed up in Figure 5b where during the 10 isometric contractions the 4 month old dystrophin deficient EDL muscles have a similar force deficit and recovery to age matched littermate controls. In contrast the 22 month old dystrophin deficient muscles with extensive complex branching (Figures 1, 9) only generate around 50% of their starting force by the 10^th^ isometric contraction, and this recovers to 60% of maximal. These maximal isometric forces are unlikely to be experienced *in vivo*, non-the-less it supports our contention that extensively branched *mdx* fast-twitch EDL muscle fibers cannot sustain maximal isometric forces due to the branch points mechanically weakening the fiber (Figures 10, 11). Claflin and Brooks ^61^ showed that 5-13 month *mdx* lumbrical muscles experience an irreversible loss of force when subjected to 10 isometric contractions, however, they did not look at fiber branching. When modeling stretching of a muscle fiber Iyer, et al. ^58^ “showed non-uniform strain distributions at branch points in single fibers, whereas uniform strain distribution was observed in fibers with normal morphology and concluded this increased susceptibility to stretch-induced damage occurring in branched myofibers at the branch point”. Further evidence that extensive branching structurally weaken the dystrophic fiber making it prone to rupture when producing high forces comes from using the skinned muscle fiber technique to study single branched fibers. When a branched dystrophic skinned fiber was attached to a force transducer, the fiber commonly breaks at a branch point when exposed to a series of solutions with increasing [Ca^2+^]. Importantly, when the remaining non-branched segment of the same fiber was reattached it could sustain maximal [Ca^2+^] activated force ^16, 51^. Additional evidence from past studies performed on merosin deficient dystrophic mice with extensive branching have shown that the branched fibers are damaged by isometric contractions ^51^.

### Reversible and irreversible recovery of force post eccentric contractions

In figure 6a-c we show that 9-22 month old *mdx* with extensive complexed branching (Figure 1, 9) have a catastrophic force deficit on the first of six ECs. The 4 month *mdx* have a graded force loss over the six ECs which, though more marked, is similar in profile to controls (Figure 6a,b). We attribute the catastrophic force drop on the first EC in older 9-22 month old *mdx* (Figure 6c) on to the structural weakness cause by the presence of large numbers of complexed branched fibers (Figure 1, 9) present in the 9-22 month old dystrophic EDL. Previously, we and others, have proposed that extensively complex branching structurally weakens fast-twitch fibers ^8, 12, 25, 50–55, 72, 73, 86–90^. In Figure 6b, both 4 months and 22 months *mdx* are dystrophin deficient, but as the animals age the absence of dystrophin triggers cycles of necrosis/regeneration and an age related increase in complex branched fibers which we propose rupture on the first EC (Figure 6b lower trace and Figure 1, 9, 11). Olthoff, et al. ^44^ showed in 3 month old *mdx* EDL muscles that there was a graded 90% force deficit which occurred over 10 EC given at 3 minutes intervals. The authors attributed the bulk of this force deficit to mechanisms linked to pathological ROS production in the dystrophic 3 month EDL muscles because there was a 65% recovery of the force within 120 minute and when EC cycles were delivered at 30 minute intervals there was a reversible loss of force. The study concludes that these experiments **<u>do not support</u>** the hypothesis that the EC force deficit in dystrophic EDL is due to sarcolemmal rupturing ^101^, as would be the case if dystrophin was acting as a shock absorber. Olthoff, et al. ^44^ hypothesized that the ECs in their EDL muscles from 3 month old mice drives a transient, redox based inhibition of contractility. Lindsay, et al. ^23^ demonstrated a similar result, again in 3 month *mdx* EDL muscles, where following 75% eccentric force deficit dystrophic muscles recovered up to 64% at 60 minutes post contraction. In support of the redox hypothesis, when Lindsay, et al. ^23^ added the macrophage synthesized antioxidant 7,8-Dihydroneopterin to the 3 month old *mdx* muscles it provided protection against ECs and improved force recovery to 81%. The authors conclude that the restoration of isometric tetanic force with antioxidant treatment in *mdx* muscle suggest reversible oxidation of proteins regulating muscle contraction. In both these cases and other studies reporting force recovery post EC in fast-twitch muscle from young *mdx* mice (3-12 weeks of age)^22, 28, 102^, according to our 2-stage model of damage in the dystrophinopathies, dystrophic fast-twitch fiber branching had not reached “tipping point” in stage 2. When the “tipping point” is passed there is an irrevocable sarcolemma rupture at branch points. In our 2018 paper we showed it was only in old *mdx* EDL with extensively branched fibers that there was a loss of ∼65% of the force on the first EC which did not recover ^12^. Here we have confirmed this finding and extended it by looking at recovery post EC at 120 minutes and by showing that in 4 months old *mdx* EDL muscles, (with considerably less branching) there is a graded force loss over the six EC which recovers by ∼50% after 120 minutes compared to only a ∼20% recovery in the old EDLs with extensively complexed branching. It should be noted that our EC protocol is severe enough to produce a non-recoverable force deficit of 20-30% in age matched littermate controls. Our current findings along with the review by Allen, et al. ^36^ and recent work mentioned ^23, 44, 45^ add to the growing body of evidence in support of our 2-stage hypothesis to explain the pathology of the dystrophinopathies ^12, 15–17, 25^.

### Summary

In the present study we have demonstrated that *mdx* EDL muscles from adult to senescent age groups have the largest component of eccentric damage which occurs as an abrupt loss of force on the first EC (Figure 6a-c). As previously reported we propose this is the result of mechanical rupture of branched dystrophic fibers ^10, 12^. In contrast younger adolescent *mdx* muscle have a graded drop in force during each phase of the EC protocol ^28, 80^, resembling those seen in littermate control muscles (Figure 6a,b). When dystrophic muscles are left to recover 120 minutes post EC we also report rapidly reversible recovery of EC force deficit in young adolescent animals but to a lesser degree (∼50% c.f. ∼65% in Olthoff, et al. ^44^) than those published previously (Figure 7c,d). Given Lindsay, et al. ^45^ report that the EC strain is the major predictor of the force deficit, differences in the present study are likely attributable to the increased magnitude in EC strain we used (Figure 8b). In aged and senescent dystrophic animals recovery is only ∼20-30% of starting force (Figure 7), indicating a smaller reversible force loss component and a larger irreversible component due to acute membrane rupture. We attribute this irreversible component to the presence of complex branches in aged and senescent *mdx* mice which are prone to rupture following EC induced injury becoming functionally obsolete. Dependent on the EC strain, degree of branching and complexity of branching the capacity for recovery varies in *mdx* muscle and can explain the variability reported in literature ^8, 9, 21–23, 28, 44, 80, 102^. Our current findings highlight the importance of studying the muscle pathophysiology in the *mdx* dystrophin-deficient mouse at all age points throughout the dystrophic animals’ life span. Many studies report skeletal muscle pathologies in *mdx* mice between 6 to 12 weeks of age yet fail to address why these dystrophin-deficient animals can live largely asymptomatically past 108 weeks of age. It is only in the old *mdx* mouse that we start to see them die earlier than age matched controls ^46, 47^ and we correlate this with the degree and complexity of muscle fiber branching where a significant proportion of branches can no longer withstand the normal stresses and strains of muscle contraction.

